# FibrilNet maps conserved and tissue-specific molecular environments across systemic amyloidoses

**DOI:** 10.64898/2026.08.29.747993

**Authors:** Pietro H. Guzzi, Ugo Lomoio, Valentina Carbonari, Pietro Lió, Pierangelo Veltri

## Abstract

Systemic amyloidoses are initiated by distinct amyloidogenic precursor proteins but frequently contain recurrent extracellular, complement, lipid-transport and matrix-remodelling components. Whether these recurrent proteins form a conserved systems-level environment across amyloid diseases, and how strongly that environment depends on precursor and tissue context, remains unresolved. We developed **FibrilNet**, a network framework that integrates experimentally defined amyloid proteomes with a human protein–protein interaction graph and Gene Ontology-derived semantic information. FibrilNet compares topology-only random walk with restart (RWR) with ontology-aware semantic-RWR in frozen leave-one-out module reconstruction and precursor-seeded prioritization tasks. The human graph contains 17,997 proteins and 925,977 physical interactions, with a 9-dimensional semantic representation of interaction context. In expanded cardiac transthyretin amyloidosis (ATTR), semantic-RWR increased mean reciprocal rank (MRR) from 0.00167 to 0.05015 and Recall@100 from 0.0199 to 0.3377, improving 132 of 151 held-out targets. Significant semantic gains were also observed in renal serum amyloid A amyloidosis (AA) and leukocyte chemotactic factor 2 amyloidosis (ALECT2). Across compact ATTR, light-chain amyloidosis (AL), AA and ALECT2 modules, APCS, VTN and TIMP3 formed a direct four-disease recurrent core, while APOE occurred in three of four modules. A tissue-aware ATTR analysis showed limited overlap between cardiac and neurologic modules (19 shared proteins; Jaccard 0.0569). In the hTTR-A97S peripheral-nerve model, semantic-RWR significantly improved reconstruction of the 202-protein mapped neurologic module, with the strongest evidence concentrated in the downregulated proteomic program. TTR-seeded propagation improved with semantic information but remained weak in absolute terms, separating precursor identity from the distributed downstream molecular environment. These results support a multilayer model in which a restricted conserved amyloid environment coexists with precursor-, tissue- and disease-specific organization.

## 1 Introduction

Amyloidoses are protein-misfolding disorders in which normally soluble proteins undergo conformational conversion and accumulate as insoluble fibrillar deposits. Systemic amyloidoses are unusually informative from a systems-biology perspective because distinct precursors can generate partially convergent tissue pathology. Transthyretin amyloidosis (ATTR), immunoglobulin light-chain amyloidosis (AL), serum amyloid A amyloidosis (AA), leukocyte chemotactic factor 2 amyloidosis (ALECT2), fibrinogen A*α* amyloidosis and apolipoprotein-derived forms differ in molecular origin, genetics and organ tropism, yet amyloid deposits recurrently contain non-fibrillar proteins involved in extracellular matrix organization, complement, lipid transport and proteostasis. Proteomic studies have directly exposed this molecular complexity. In cardiac ATTR and AL, laser-microdissection mass spectrometry identified both a broad expanded plaque proteome and a smaller amyloid-specific proteome, with shared and disease-specific components [1]. A large kidney amyloid atlas subsequently demonstrated extensive between- and within-disease heterogeneity across renal amyloid types, including AA and ALECT2 [2]. These datasets suggest that amyloid deposition should be viewed not only as accumulation of a precursor protein but also as the emergence of a distributed molecular environment.

A network formulation is attractive because proteins participating in the same pathological environment need not be directly adjacent to the precursor. Random-walk methods have long been used to prioritize disease genes from interaction networks [9], but conventional diffusion treats network edges as functionally interchangeable. Protein–protein interaction graphs are context-agnostic: two edges may both be physical interactions while participating in very different biological processes. We therefore asked whether ontology-derived functional information could provide a biologically meaningful inductive bias for information propagation.

We developed **FibrilNet**, which combines a human protein–protein interaction network with Gene Ontology (GO)-derived semantic features. The interaction graph is built from curated human physical interactions and mapped to reviewed proteins; GO annotations provide biological process, molecular function and cellular component context [6, 7]. FibrilNet evaluates topology-only RWR against semantic-RWR, in which propagation is biased toward interactions with greater ontology compatibility. Importantly, the disease analyses use frozen graph context and experimentally defined protein modules; disease labels are not used to retrain the underlying representation.

Our hypothesis is that distinct amyloidogenic precursor proteins may converge on a restricted conserved deposition environment while retaining precursor-, tissue- and disease-specific pathological modules. We therefore represent each disease environment as

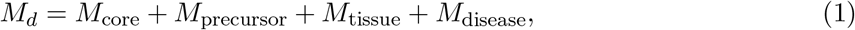

and refine ATTR as

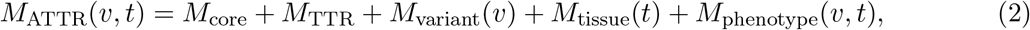

where *v* denotes TTR variant and *t* tissue context. ATTR is particularly suitable for this analysis because inherited variants can be predominantly neurologic, predominantly cardiac, or mixed, with phenotype additionally influenced by age and geography [5].

We address four questions: (i) whether semantic diffusion improves reconstruction of experimentally defined amyloid modules; (ii) whether independent amyloidoses share a recurrent molecular core; (iii) whether semantic organization differs by disease and tissue; and (iv) whether the amyloidogenic precursor alone is sufficient to recover the broader downstream environment. The overall workflow is shown in fig. 1.

**Figure 1.**
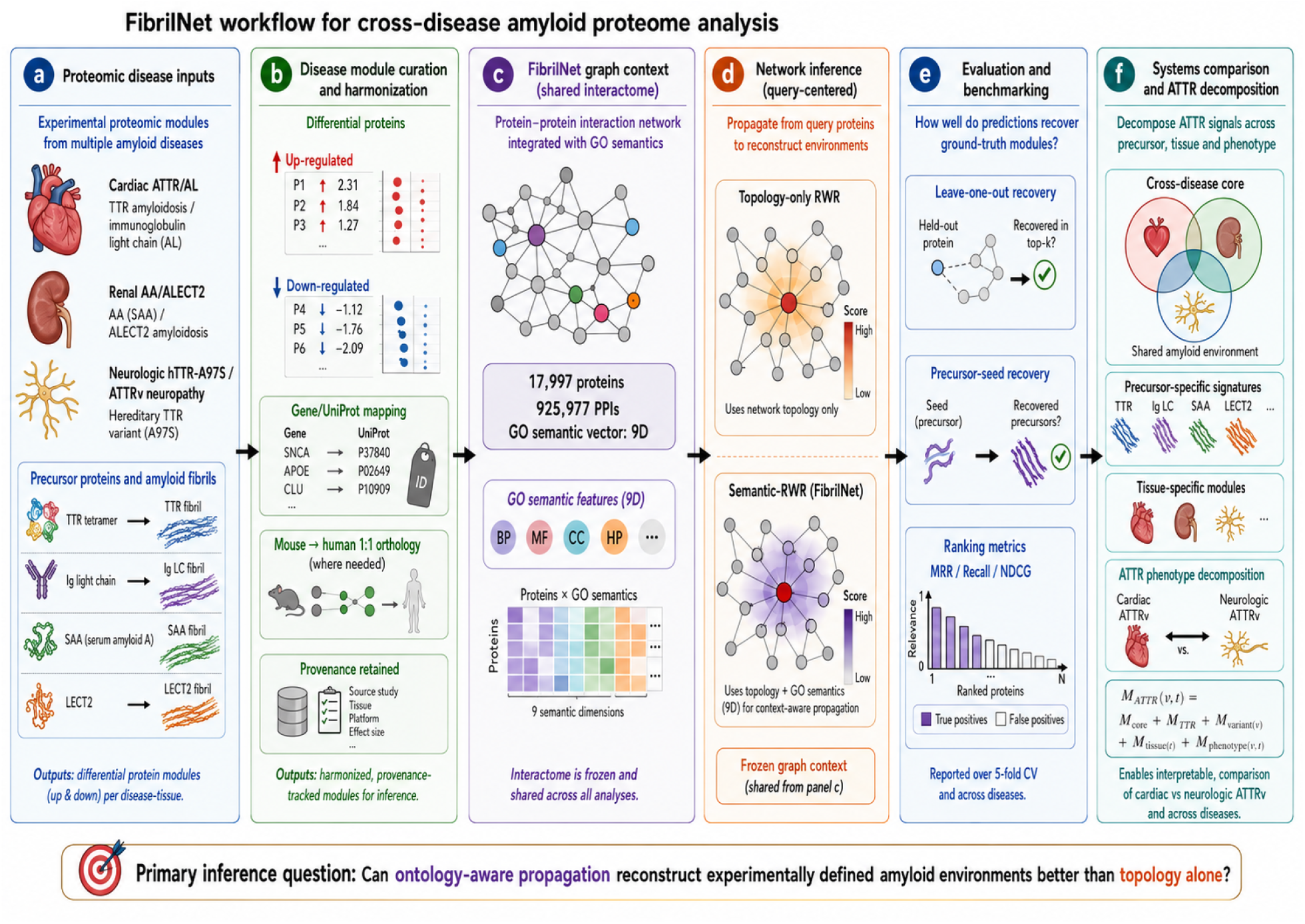
FibrilNet workflow and tissue-aware ATTR model. Experimentally defined proteomic modules are harmonized to the human FibrilNet graph, followed by topology-only and ontology-aware diffusion. Primary evaluation uses symmetric leave-one-out module completion; secondary analyses use precursor-seeded recovery, cross-disease overlap and tissue/direction stratification. The human graph used here contains 17,997 proteins, 925,977 physical interactions and 9-dimensional GO-derived semantic edge information.

## 2 Results

### 2.1 Cardiac ATTR and AL share a plaque environment but retain disease-specific structure

We first analyzed the amyloid-specific cardiac proteomes reported by Kourelis et al. [1]. Twenty-seven ATTR proteins and 16 AL proteins mapped to the FibrilNet graph, with 12 shared proteins (Jaccard = 0.387). This direct overlap was substantially greater than cross-tissue overlap involving the renal modules, consistent with a strong contribution of organ context to the observed amyloid environment.

In symmetric leave-one-out module completion, semantic-RWR improved 26 of 27 ATTR targets and 15 of 16 AL targets. For ATTR, semantic-RWR achieved MRR = 0.0937 and Recall@100 = 0.370, with a one-sided paired Wilcoxon *p* = 1.49 *×* 10^−8^. For AL, semantic-RWR achieved MRR = 0.0371 and Recall@100 = 0.250 (*p* = 3.05 *×* 10^−5^). The result indicates that ontology-aware paths provide information beyond graph topology in two independent cardiac amyloid types.

### 2.2 Expanded cardiac ATTR shows strong distributed semantic coherence

We next tested whether the effect generalized beyond the compact amyloid-specific ATTR signature. The published expanded cardiac ATTR proteome contained 160 gene-level proteins [1]; 151 mapped uniquely to the human graph. The source table is a membership list rather than a ranked evidence list, so robustness was evaluated by repeated random subsets of 20, 30, 50 and 100 proteins in addition to the full module.

For the complete 151-protein module, topology-only RWR achieved MRR = 0.00167, median rank = 1684, Recall@100 = 0.0199 and Recall@200 = 0.0596. Semantic-RWR increased MRR to 0.05015, improved median rank to 289, and raised Recall@100 and Recall@200 to 0.3377 and 0.4238, respectively. Semantic-RWR improved 132 of 151 targets, with a median semantic/topological rank ratio of 0.253 and a one-sided Wilcoxon *p* = 4.46 *×* 10^−23^.

The effect was robust to module size. All replicates were positive at sizes 30, 50 and 100, while 19 of 20 size-20 replicates showed positive ΔMRR. This sensitivity analysis argues that the result reflects distributed organization of the expanded plaque environment rather than dependence on a small fixed signature (Supplementary Fig. S1).

### 2.3 TTR alone does not reconstruct the cardiac plaque environment

We then contrasted distributed module reconstruction with precursor-centred propagation from TTR. In the original amyloid-specific ATTR module, TTR-seeded RWR produced low early-rank recovery; semantic-RWR modestly improved overall ordering, especially among shared plaque proteins, but did not recover the ATTR-specific submodule at early ranks. Thus, the strong reconstructability of the distributed ATTR module does not imply that plaque proteins form a local neighbourhood around TTR.

This distinction supports a two-level organization: TTR is the causal amyloidogenic precursor, while the downstream plaque environment is a distributed network state whose organization becomes visible only when multiple disease-associated proteins provide context.

### 2.4 Renal AA and ALECT2 generalize semantic reconstruction beyond cardiac amyloidosis

We next analyzed the kidney amyloid proteomic atlas of Charalampous et al. [2]. The AA contrast comprised 418 AA cases against 2,130 comparator amyloid cases across 496 measured protein columns. After two-sided Mann–Whitney testing, Benjamini–Hochberg correction and positive-effect filtering, 59 proteins were significantly enriched and 48 mapped into the complete primary graph module.

For the complete mapped AA module, RWR achieved MRR = 0.00249, whereas semantic-RWR reached 0.09061. Median rank improved from 2201.5 to 783.5, Recall@100 increased from 0.0625 to 0.2292, and semantic-RWR improved 42 of 48 targets. The paired ΔMRR was 0.0881 (95% bootstrap CI 0.0282–0.1641), with one-sided Wilcoxon *p* = 2.04 *×* 10^−10^. Positive effects were retained across top-10, top-20, top-30 and top-50 sensitivity tiers (Supplementary Fig. S2).

ALECT2 was analyzed against immunoglobulin amyloidosis within the same renal atlas, providing an organ-matched comparison. The contrast included 474 ALECT2 and 1,618 AL/AH cases, with 143 positively significant mapped candidates and 124 proteins in the primary mapped module. Semantic-RWR improved 100 of 124 held-out targets. MRR increased from 0.00239 to 0.01614, median rank improved from 1855 to 837, and Recall@100 increased from 0.0242 to 0.2419. The paired improvement was highly significant (*p* = 1.06 *×* 10^−13^), with ΔMRR = 0.01375 (95% CI 0.00439–0.02586).

By contrast, propagation from the LECT2 precursor alone did not improve global MRR, providing a second example in which a causal amyloidogenic precursor and a distributed downstream environment are not equivalent network objects.

### 2.5 A restricted molecular core recurs across four systemic amyloidoses

We compared compact ATTR, AL, AA and ALECT2 modules directly. Pairwise overlap was highest for cardiac ATTR–AL (Jaccard = 0.387) and substantially lower across cardiac–renal comparisons (Jaccard 0.067–0.136; fig. 2C). We defined the direct feature-enrichment core frequency

**Figure 2.**
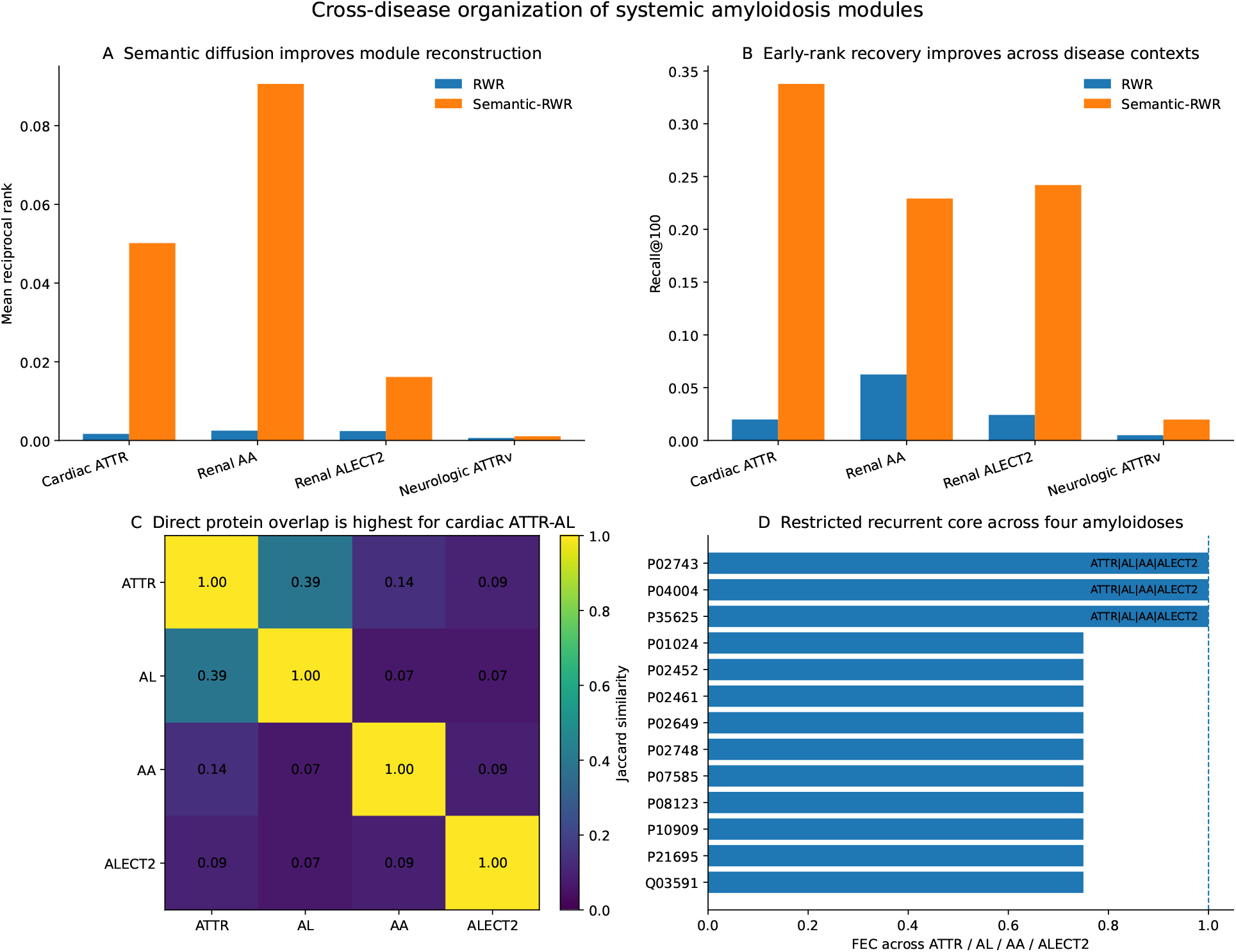
Cross-disease organization of systemic amyloidosis modules. **A**, Mean reciprocal rank for topology-only RWR and semantic-RWR in expanded cardiac ATTR, renal AA, renal ALECT2 and the neurologic ATTRv module. **B**, Corresponding Recall@100. **C**, Pairwise Jaccard overlap among compact ATTR, AL, AA and ALECT2 modules. **D**, Highest direct FEC values across the four disease modules; APCS (P02743), VTN (P04004) and TIMP3 (P35625) occur in all four. Absolute metric values should be interpreted within module because module size and experimental source differ across diseases.

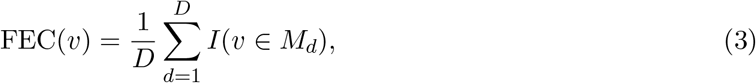

where *D* = 4 disease modules.

Three proteins—APCS, VTN and TIMP3—were present in all four modules (FEC = 1), defining the strict direct core. APOE was present in ATTR, AL and AA (FEC = 0.75). Several complement and matrix-associated proteins also recurred in three of four diseases. These recurrent proteins are not interpreted as universal causal drivers; rather, they identify points of cross-disease convergence in experimentally observed amyloid environments.

### 2.6 Neurologic ATTRv is largely distinct from cardiac ATTR and shows weaker, direction-specific semantic organization

ATTR is a systemic disease with genotype- and tissue-dependent phenotypes rather than a purely cardiac disorder [5]. We therefore extended FibrilNet to a neurologic context using the sural-nerve proteome of the hTTR-A97S knock-in model from PXD054291 [3]. The underlying study identified early cytoskeletal abnormalities preceding overt axonal degeneration. Human clinical skin-biopsy data independently support amyloid deposition and small-fibre loss as relevant markers of ATTRv neuropathy [4].

The source proteomic analysis contained 245 differential protein rows, comprising 46 increased and 199 decreased proteins. Mouse protein accessions were resolved to gene symbols and mapped to explicit human one-to-one Ensembl orthologs. This produced 213 unique humanized proteins (40 upregulated and 173 downregulated); 202 mapped to the FibrilNet graph.

Direct overlap with the expanded cardiac ATTR module was low: 19 proteins were shared between 151 cardiac and 202 neurologic mapped proteins (Jaccard = 0.0569; fig. 3A). Thus, the two ATTR environments are dominated by tissue-specific components despite sharing TTR as the amyloidogenic precursor.

**Figure 3.**
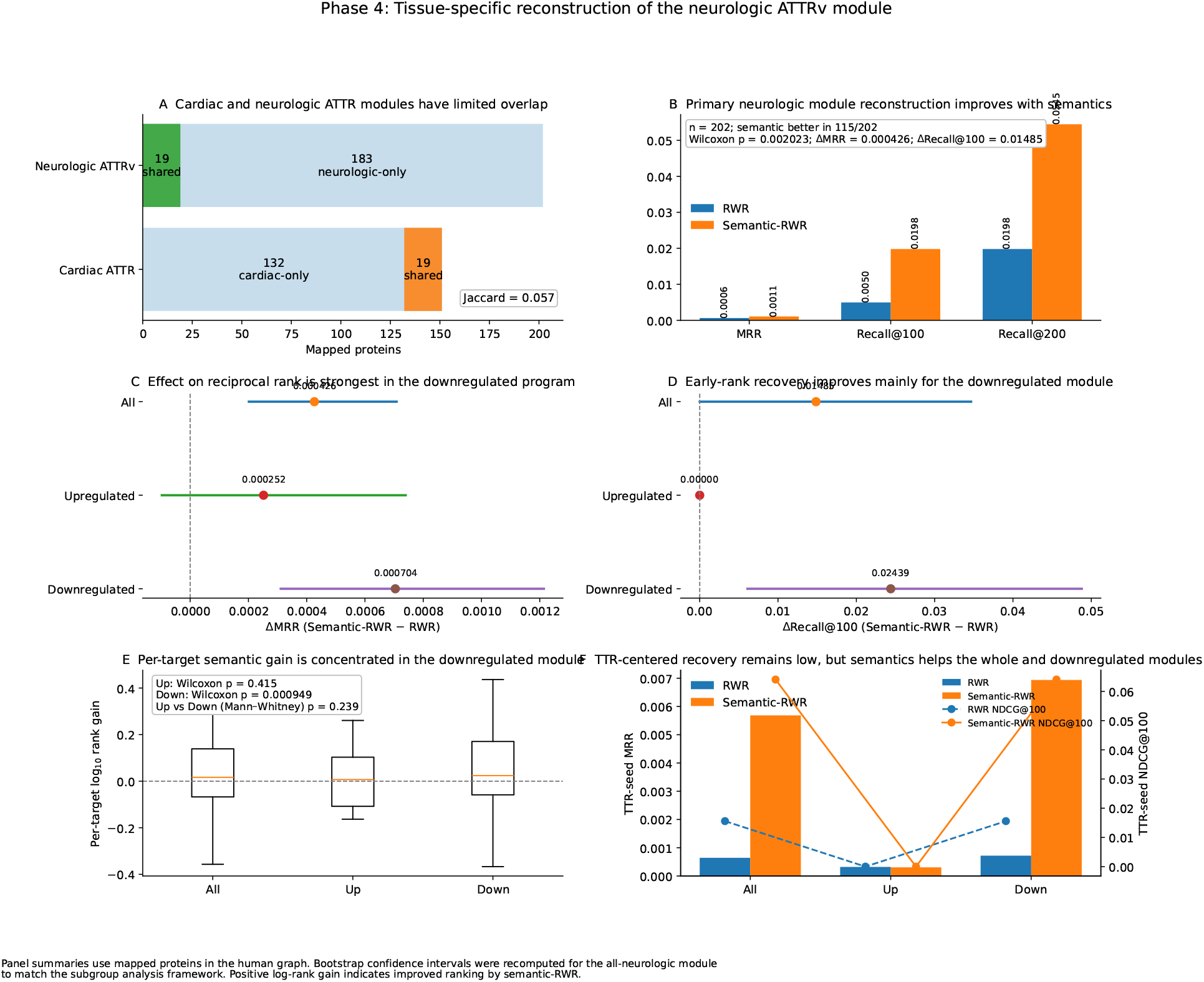
Phase 4 reveals a tissue-specific and predominantly downregulated neurologic ATTRv module. **A**, Cardiac versus neurologic ATTR overlap. **B**, Primary neurologic leave-one-out reconstruction. **C–D**, Bootstrap effects for MRR and Recall@100 in the complete, upregulated and downregulated modules. **E**, Per-target semantic rank-gain distributions. **F**, TTR-seeded recovery for the complete and direction-stratified neurologic modules. Positive semantic effects are strongest and statistically robust in the downregulated program, whereas direct up-versus-down gain differences are not significant.

In the primary 202-protein neurologic leave-one-out task, semantic-RWR improved MRR from 0.000630 to 0.001056, median rank from 4458.5 to 4306.5, Recall@100 from 0.00495 to 0.01980, and Recall@200 from 0.01980 to 0.05446. Semantic-RWR improved 115 of 202 targets and the paired effect was significant (*p* = 0.0020).

Direction stratification showed that this signal was concentrated in the downregulated program. Among 38 mapped upregulated proteins, the semantic improvement was not significant after Holm correction (*p* = 0.415), and the bootstrap interval for ΔMRR crossed zero. Among 164 mapped downregulated proteins, semantic-RWR increased MRR from 0.000719 to 0.001423, improved median rank from 4104.5 to 3862.5, raised Recall@100 from 0.0122 to 0.0366, and improved 98 of 164 targets. The Holm-adjusted Wilcoxon *p* was 9.49 *×* 10^−4^; ΔMRR was 0.000704 (95% CI 0.000311–0.001214) and ΔRecall@100 was 0.02439 (95% CI 0.00610–0.04878). The direct difference in per-target semantic gain between upregulated and downregulated groups did not reach significance (Mann–Whitney *p* = 0.239), so the asymmetry is interpreted conservatively.

### 2.7 TTR-centred semantic propagation improves focus but not absolute recovery of the neurologic environment

Finally, we evaluated whether TTR alone could recover the neurologic module. Topology-only TTR-seeded RWR achieved MRR = 0.000643, Recall@100 = 0.00995 and NDCG@100 = 0.0156. Semantic-RWR improved these values to MRR = 0.005684, Recall@100 = 0.01493 and NDCG@100 = 0.0641. The improvement was concentrated in the downregulated subgroup, where semantic-RWR increased TTR-seeded MRR from 0.000718 to 0.006938. The upregulated subgroup showed essentially no early TTR-seeded recovery by either method.

Thus, semantic information makes TTR-centred propagation more biologically focused, but absolute recovery remains low. This result parallels the cardiac analysis and supports separation of the precursor layer from the downstream tissue-associated network state.

## 3 Discussion

FibrilNet provides a systems-level interpretation of systemic amyloidosis proteomes. The central observation is not that all amyloidoses share one molecular module, but that a restricted recurrent environment coexists with strong disease- and tissue-specific organization. Across cardiac ATTR and AL and renal AA and ALECT2, ontology-aware diffusion consistently improved reconstruction of experimentally defined modules relative to topology alone. The magnitude of the effect varied markedly, indicating that semantic organization is itself a property of disease context rather than a fixed feature of the interactome.

The strongest effect occurred in the expanded cardiac ATTR plaque environment. Semantic-RWR improved early-rank recovery by more than an order of magnitude and remained robust under repeated random subsampling. This result is consistent with the cardiac plaque proteome containing a distributed but functionally coherent extracellular and host-response environment. The compact ATTR and AL modules also showed substantial direct overlap, in agreement with the original proteomic evidence that the two cardiac amyloid types contain both common and distinct components [1].

The renal analyses generalize the result across precursor proteins and an independent organ. Both AA and ALECT2 showed significant semantic gains across multiple sensitivity tiers. Because AA and ALECT2 were derived from the same renal atlas, their contrast reduces one source of experimental heterogeneity compared with cross-organ comparisons. At the same time, the strict direct overlap among ATTR, AL, AA and ALECT2 was small. APCS, VTN and TIMP3 formed the only FEC=1 core, with APOE present in three of four modules. We interpret this as evidence for restricted convergence rather than a large universal amyloid signature.

The ATTR analyses provide a second conceptual result: precursor identity and downstream tissue organization are partially separable. TTR is the common causal precursor in both cardiac and neurologic ATTR, yet cardiac and peripheral-nerve modules showed only 19 shared mapped proteins. Moreover, distributed cardiac ATTR showed very strong semantic coherence, while the neurologic module showed a smaller effect concentrated in proteins decreased in hTTR-A97S nerve. This difference is compatible with tissue-specific network responses superimposed on a common precursor.

The direction-stratified neurologic result is particularly relevant to the biology of the hTTR-A97S model. The source study identified cytoskeletal abnormalities and axonal dysfunction preceding overt degeneration [3]. FibrilNet does not establish that downregulation causes neuropathy, but the stronger semantic organization of the downregulated program is consistent with coordinated disruption of established neuronal and axonal functions. The lack of a significant direct difference in gain distributions between up- and downregulated groups prevents a categorical mechanistic interpretation, and the subgroup result should therefore be considered supportive rather than definitive.

Precursor-seeded analyses reinforce the multilayer view. TTR-centred semantic propagation improved ranking relative to topology, but neither cardiac nor neurologic environments were strongly recoverable from TTR alone. Likewise, the LECT2 seed did not globally reconstruct the ALECT2 module. These observations do not challenge the causal role of the precursor proteins. Instead, they indicate that the mature amyloid-associated environment is a distributed network state that cannot be reduced to local graph proximity around the fibril-forming protein.

This distinction motivates the ATTR decomposition

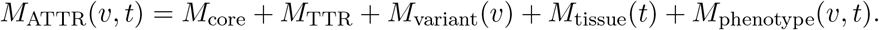

The present data directly support separation of the shared precursor from tissue-associated down-stream organization. The variant and phenotype terms remain to be estimated from genuine variant-resolved human molecular datasets. This is important because ATTRv can manifest predominantly neurologic, predominantly cardiac or mixed phenotypes [5]; synthetic assignment of variant-specific proteomes would therefore be inappropriate.

From a methodological perspective, FibrilNet illustrates a general strategy for network medicine: ontology-derived information can be used to bias propagation through a context-agnostic interactome without replacing physical interactions. The approach is deliberately conservative. It does not infer new physical interactions, and it does not treat semantic proximity as causality. Rather, semantic-RWR asks whether known disease-associated proteins are more recoverable when graph information flow is preferentially routed through functionally compatible interactions.

## 4 Methods

### 4.1 Human interaction graph

The FibrilNet graph was constructed from curated human physical interactions, mapped to reviewed human proteins. The benchmark graph contains 17,997 nodes and 925,977 unique physical protein– protein interactions. Gene Ontology annotations were used to encode semantic context across biological process, molecular function and cellular component [6]. The underlying protein-interaction resource was BioGRID [7]; curated complex information was retained as an independent benchmark resource using Complex Portal [8].

For each interaction, the semantic representation contains three values for each GO branch: information-content-based node similarity, information-content-based path similarity and an annotation-availability mask, yielding a 9-dimensional semantic vector.

### 4.2 Topology-only and semantic random walk with restart

For an undirected graph with transition matrix *P*, topology-only RWR iterates

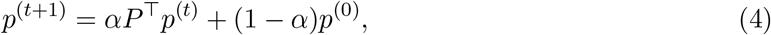

where *p*^(0)^ is concentrated on the query set and *α* is the continuation probability. Semantic-RWR uses an alternative transition matrix in which interaction probabilities are modulated by semantic compatibility before row normalization. The same restart parameter and query definitions are used for both methods, so the comparison isolates the effect of ontology-aware propagation.

### 4.3 Leave-one-out disease-module reconstruction

For disease module *M*_*d*_ and target *v* ∈ *M*_*d*_, the query set is

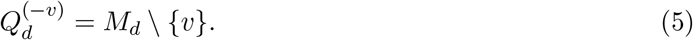

The target is removed from the query set, RWR and semantic-RWR are run from the remaining module, and all graph candidates are ranked. Primary metrics are MRR and Recall@20, @50, @100 and @200. This task evaluates recoverability of experimentally defined modules without disease-label retraining.

### 4.4 Cardiac ATTR and AL

Cardiac modules were derived from the proteomic atlas of Kourelis et al. [1]. The compact amyloid-specific analysis used mapped ATTR and AL signatures with symmetric leave-one-out evaluation. The expanded ATTR analysis used the published 160-protein non-amyloidogenic plaque proteome after gene-level normalization; 151 proteins mapped uniquely to FibrilNet. Because the published list defines membership rather than rank, robustness was tested by 20 random replicates at module sizes 20, 30, 50 and 100, plus the complete mapped module.

### 4.5 Renal AA and ALECT2

Renal modules were derived from the processed kidney amyloid proteome of Charalampous et al. [2]. For AA, cases were contrasted against the comparator amyloid group after excluding AA, AFib and AApoCII from the comparator definition used in the analysis. For ALECT2, 474 ALECT2 cases were contrasted with 1,618 immunoglobulin amyloidosis cases. Each protein column was tested using a two-sided Mann–Whitney U test, followed by Benjamini–Hochberg correction. Effect size was the log_2_ ratio of mean normalized spectral counts using a half-minimum-positive pseudocount. Primary modules required FDR *<* 0.05 and positive effect. Sensitivity tiers used the top 10, 20, 30 and 50 positively significant mapped proteins plus the complete mapped module.

### 4.6 Neurologic ATTRv module and orthology

The neurologic molecular source was PXD054291, associated with the hTTR-A97S sural-nerve study [3]. The publication-defined filtered proteomic set contained 245 differential proteins (46 increased and 199 decreased). Protein accessions were resolved to mouse gene symbols through UniProt when necessary. Mouse genes were then mapped to human orthologs through explicit Ensembl homology, and only one-to-one orthologs were retained for the primary human-network analysis. This yielded 213 unique humanized proteins, including 40 upregulated and 173 downregulated proteins; 202 mapped to FibrilNet.

The complete mapped neurologic module was the prespecified primary analysis. Upregulated and downregulated modules were secondary strata. For the two stratified Wilcoxon tests, Holm correction was applied. Paired bootstrap confidence intervals used 20,000 resamples. A Mann– Whitney test compared per-target log_10_ rank gain between up- and downregulated strata as an exploratory analysis.

### 4.7 Precursor-seeded analyses

TTR and LECT2 were evaluated separately from distributed module completion. A precursor node was used as the sole query and the corresponding disease module was treated as the positive set. These experiments test whether the precursor’s graph neighbourhood is sufficient to recover the broader molecular environment. They are network-recovery analyses and should not be interpreted as tests of biochemical causality.

### 4.8 Cross-disease overlap and recurrent core

For disease modules *A* and *B*, direct overlap was quantified by Jaccard similarity

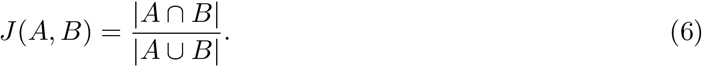

For protein *v* across *D* modules, FEC was defined as

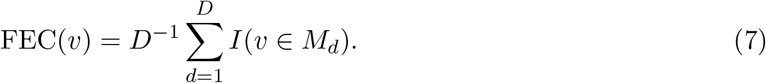

Cross-tissue comparisons are interpreted separately from tissue-matched comparisons because organ, species and proteomic design can all contribute to observed differences.

### 4.9 Statistical analysis

Paired target-rank improvements were tested using one-sided Wilcoxon signed-rank tests where the prespecified alternative was improved semantic ranking. Bootstrap confidence intervals were calculated by paired resampling of targets. For the neurologic directional analysis, Holm correction was applied across the two prespecified subgroup tests. All exact module-level metrics were generated by the released FibrilNet scripts.

## 5 Limitations

The study has several limitations. First, the modules were derived from different tissues and experimental designs. Cardiac ATTR and AL were measured in human plaques, AA and ALECT2 were derived from a human renal atlas, and the neurologic ATTRv analysis used a mouse sural-nerve model followed by human orthology mapping. Absolute performance values should therefore not be compared as if module size, tissue and sampling were identical.

Second, the neurologic analysis is a cross-species mechanistic comparison rather than direct human cardiac-versus-human nerve proteomics. One-to-one orthology increases mapping specificity but does not guarantee conservation of expression regulation or tissue context. Human variant- and tissue-resolved ATTRv molecular datasets are required to estimate variant-specific effects directly.

Third, protein-interaction networks are incomplete and literature-biased, and GO annotation depth is non-uniform. Semantic-RWR can reduce the influence of functionally implausible paths but cannot eliminate missing biology or annotation bias.

Fourth, direct FEC depends on module definition and source coverage. APCS, VTN and TIMP3 are a strict core of the four analyzed modules, not a claim of a universal amyloid core across every hereditary or acquired amyloidosis.

Finally, precursor-seeded recoverability is not equivalent to biochemical initiation. Weak TTR- or LECT2-seed performance does not diminish the established causal role of those amyloidogenic proteins; it shows that the broader disease environment is not represented by precursor-centred graph proximity alone.

## 6 Conclusion

FibrilNet identifies a reproducible systems-level organization across systemic amyloidoses. Ontology-aware diffusion improves reconstruction of experimentally defined cardiac and renal amyloid environments, while a restricted cross-disease core coexists with strong precursor- and tissue-specific structure. The expanded cardiac ATTR environment is particularly coherent under semantic propagation. In neurologic ATTRv, direct overlap with the cardiac environment is low and the strongest semantic organization is found in the broad downregulated peripheral-nerve program. Across both cardiac and neurologic ATTR, precursor-seeded recovery remains limited, supporting a multilayer architecture in which the amyloidogenic precursor and the downstream tissue-associated molecular environment are partially separable. FibrilNet provides a reproducible framework for extending this analysis to additional systemic amyloidoses and future human variant-resolved datasets.

## Data availability

The study reuses publicly available proteomic resources: the cardiac amyloid plaque atlas [1], the kidney amyloid proteomic atlas [2], and PXD054291 [3]. Derived disease modules, mapping tables, analysis outputs and figure source data are included in the accompanying FibrilNet archival package. The neurologic source data remain traceable to PXD054291.

## Code availability

The complete FibrilNet analysis code, frozen derived inputs, result tables, environment specifications and reproduction scripts are archived on Zenodo at 10.5281/zenodo.22161436.

## Author contributions

All the authors conceived the study, designed the computational framework, performed the analyses, interpreted the results and wrote the manuscript.

